# Bacterial communities of premise plumbing systems in four European cities, and their association with culturable *Legionella*

**DOI:** 10.1101/2022.08.12.503735

**Authors:** Maria Scaturro, Federica Del Chierico, Yair Motro, Angeliki Chaldoupi, Anastasia Flountzi, Jacob Moran-Gilad, Antonietta Girolamo, Thomai Koutsiomani, Bozena Krogulska, Diane Lindsay, Renata Matuszewska, Georgios Papageorgiou, Katarzyna Pancer, Nikolaos Panoussis, Maria Cristina Rota, Søren Anker Uldum, Emmanuel Velonakis, Dominique Louise Chaput, Maria Luisa Ricci

**Author notes:** Both authors contributed equally to the work. To whom correspondence should be addressed.

## Abstract

*Legionella* species are Gram negative, facultative, intracellular bacteria found in natural and engineered water systems. Understanding the bacterial interactions underlying the success of *Legionella* in aquatic environments could be beneficial for control. We aimed to profile, by 16S rRNA amplicon sequencing, the bacterial communities in premise plumbing systems of buildings in four European cities (Copenhagen, Warsaw, Rome, Athens), and identify positive and negative associations of specific community members to culturable *Legionella*. The coarse taxonomic composition was similar across the four cities, but Copenhagen and Warsaw had richer, more diverse communities than Athens and Rome, with a greater number of city-specific amplicon sequence variants (ASVs). The cities had statistically significant differences in bacterial communities at the ASV level, with relatively few shared ASVs. Out of 5,128 ASVs, 73 were classified as *Legionella*, and one or more of these were detected in most samples from each city (88.1% overall). Interestingly, the relative abundance of *Legionella* ASVs did not correlate with *Legionella* culture status. Overall, 44.2% of samples were *Legionella* culture positive: 71.4% in Warsaw, 62.2% in Athens, 22.2% in Rome, and 15.2% in Copenhagen. 54 specific ASVs and 42 genera had significant positive or negative associations with culturable *Legionella*. Negative associations included *Staphylococcus, Pseudomonas*, and *Acinetobacter*. Positive associations included several *Nitrospira* ASVs and one classified as *Nitrosomodaceae* oc32, ASVs in the amoeba-associated genera *Craurococcus-Caldovatus* and *Reyranella*, and the predatory genus *Bdellovibrio*. Some of these associations are well supported by laboratory studies, but others are the opposite of what was expected. This highlights the difficulties in translating pure culture results to into complex real-life scenarios. However, these positive and negative associations held across the four cities, across multiple buildings and plumbing compartments. This is important because developing better control measures, including probiotic approaches, will require an understanding of ecological relationships that can be generalised across different engineered water systems.

**Importance:** This study provides a snapshot of the diversity of microbial communities among premise plumbing systems in four European cities, providing new information on bacterial ASVs and genera that have positive or negative associations with culturable *Legionella* across a broad geographical and climatic range. This could inform studies aimed at confirming both *in vitro* and real-life scenarios around the role of other microbial community members in modulating *Legionella* proliferation. It could also help in the development of probiotic approaches to controlling this opportunistic pathogen.

## Introduction

*Legionella pneumophila* (Lp) can cause Legionnaires’ disease, a severe pneumonic illness that, outside the covid-19 pandemic period, has shown a growing trend in cases across European countries (www.ecdc.europa.eu/en/legionnaires-disease/surveillance/atlas). The disease is spread primarily by inhalation of *Legionella*-contaminated aerosols produced by inadequate maintenance of water systems. *Legionella* species are aquatic microorganisms often present at low concentrations in freshwater environments, but through municipal water dissemination, they can reach artificial water systems, where they often proliferate thanks to warmer temperatures and nutrient availability. *Legionella* can live in a planktonic form; but owes its survival to multi-species biofilms and protozoa from which it derives nutrients and protection from physical stress and disinfection procedures (1–3). *Legionella* abundance is modulated by complex microbial interactions, but even where its relative abundance in a biofilm is low, it can form a significant proportion of the bulk water microbiome through lysis of host amoebal cells and release of *Legionella* (4).

The survival of Lp in engineered water systems is therefore driven not only by favorable temperature and nutrient conditions but also by interactions with other microbial species found as planktonic cells or in mixed community biofilms. Such is the importance of these other community members, that a probiotic approach, one focused on encouraging competitors, grazers, and antagonists in engineered water systems, has been suggested to control proliferation of Lp (5, 6). Numerous bacterial taxa have been shown in pure culture to have a positive or negative influence on Lp growth. For example, the presence in plumbing systems of *Flavobacterium breve*, a L-cysteine producing bacterium, was shown to support Lp growth, as Lp cannot produce this amino acid itself (7). Conversely, several taxa are known to inhibit Lp. *Staphylococcus warneri* produces a peptide called warnericin that has a *Legionella*- specific antagonistic effect (8). *Pseudomonas* species can inhibit Lp through several mechanisms, including killing host *Acanthamoeba castellanii* (9), and production of biosurfactants (10), toxoflavin (11), bacteriocin-like substances (12), and volatile organic compounds with long-range inhibition (13). *Bdellovibrio*, an aerobic Gram- negative bacterium known to prey on other Gram-negative taxa, has been shown to cause lysis of at least three *Legionella* species (14). *Bacillus subtilis* can produce a surfactin that inhibits both *Legionella* and its acanthamoebal host (15). *Neochlamydia* species are obligate intracellular symbionts of amoeba, and their presence blocks infection by *Legionella*, thereby disrupting its replication (16). Numerous other taxa have been shown to reduce *Legionella* growth in pure culture, including isolates from the genera *Aeromonas*, *Flavobacterium*, *Acinetobacter*, *Kluyvera*, *Rahnella*, and *Sphingobacterium* (13).

Translating culture findings to real-life environmental systems is difficult as two or three isolated species on an agar plate will not reflect the diversity in biofilms or bulk water containing hundreds, if not thousands, of additional taxa. Although they can only show correlation rather than causation. Association studies, where abundance and prevalence of other microbial community members, with different levels of *Legionella* growth, are compared, can indicate which laboratory-based, pure culture findings are most important under natural conditions (i.e. in engineered water systems), pointing to potential targets for probiotic interventions (6). A few studies have used this approach in cooling towers in Quebec, Canada (17, 18) and across the USA (19), as well as in premise plumbing of high-rise buildings in New York, USA (20). These studies used microbial community profiling by culture-independent sequencing of bacterial 16S rRNA (17, 19, 20) or eukaryotic 18S rRNA (18) to see which other community members showed positive or negative correlations with *Legionella*. These studies, correlating microbial community taxa with presence of *Legionella*, provided real-world evidence that supports some of the findings from laboratory work with pure cultures. However, the two studies that reported *Legionella*-bacteria associations at the family or genus level were both in cooling towers in North America, so it is unclear whether these associations occur across a wider range of engineered water environments. Therefore, we aimed to determine if similar relationships between *Legionella* and other specific bacterial taxa held across a broad geographical range (four European cities with key differences in water treatment approaches) and across different buildings and premise plumbing compartments, covering a wide temperature range. This required the comparison of the resident bacterial communities in buildings sampled in the four cities. Determining which *Legionella*-community interactions are broadly universal and which are limited to specific systems or conditions will help improve probiotic and other strategies for controlling the growth of *Legionella* in engineered water systems.

## Results

Of the 204 samples collected across the four cities, 176 were included in the final analyses: 45 from Athens, 45 from Rome, 44 from Copenhagen, and 42 from Warsaw. One of the excluded samples had abnormally low sequencing depth (30 reads remaining after QC), and the rest of the excluded samples fell below the minimum 10 ng DNA threshold required by the sequencing service. Temperature and pH data were collected for a subset of the samples (n=140), and their distributions show some differences among the four cities (Figure S1). However, due to missing measurements and because other potentially important physical/chemical parameters were not measured across all samples (e.g. dissolved carbon and nitrogen content, salts, disinfectant concentration), these parameters were not included as distinct terms in statistical models, but rather they were assumed to be part of what differentiated cities, buildings and compartments, and therefore were implicitly included in these model terms.

In total, 12 390 846 sequences passed QC and were grouped into 5 128 amplicon sequence variants (ASVs). Sequencing depth per sample ranged from 2 421 to 197 630, with a median depth of 66 159. All but four samples had over 10k sequences, so for analyses where subsampling to equal depth was required, these four samples were excluded in order to increase the amount of sequence data retained across the other samples.

### Main bacterial classes and genera

Across all samples, 44 bacterial phyla were represented in the 16S rRNA data, of which 10 accounted for at least 1% of sequences (Figure 1A) and 93.0% in total. *Proteobacteria* was the dominant phylum, accounting for 66.1% of all sequences and 2 298 of the 5 128 ASVs. The proteobacterial sequences that could be further resolved all grouped into the classes *Gamma-* and *Alphaproteobacteria*. *Gammaproteobacteria* accounted for 41.4% of sequences and 1 205 ASVs overall and included 39 orders, the dominant one being *Burkholderiales* with 34.4% of total 16S sequences, followed by *Xanthomonadales* and *Pseudomonadales* both at 1.1% *Gammaproteobacteria* were most abundant in all cities except Athens, where *Alphaproteobacteria* were more abundant (Figure S2). Overall, *Alphaproteobacteria* accounted for 24.7% of sequences and 1 081 ASVs, and included 25 orders, the dominant ones being *Sphingomonadales* (8.6%), *Rhizobiales* 7.4%, *Acetobacterales* (2.3%), *Caulobacterales* (2.2%), and *Reyranellales* (1.8%).

**Figure 1.**
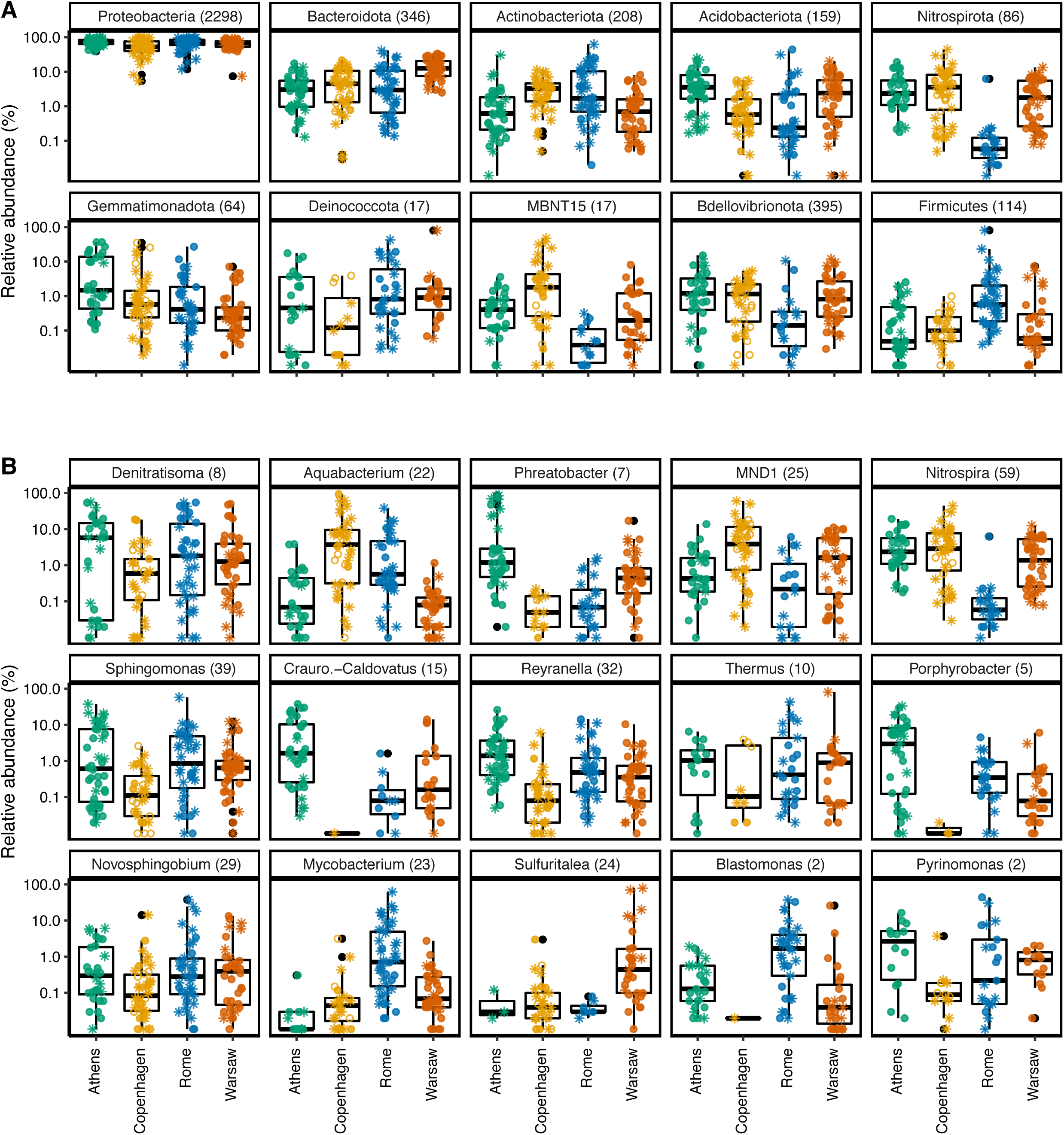
Main bacterial phyla and genera in premise plumbing systems in four European cities. (A) Top ten most abundant phyla across all samples. (B) Top fifteen most abundant genera across all samples. Parentheses next to taxon names show the number of distinct ASVs with that classification. Samples were normalised to 10 148 sequences. Filled circles show *Legionella* culture-positive samples, asterisks show culture-negative samples, open circles show samples with no culture data.

After *Proteobacteria*, the next most abundant phylum was *Bacteroidota* at 7.8% and 346 ASVs, followed by *Actinobacteriota*, *Acidobacteriota* and *Nitrospirota* at 3.5, 3.3 and 3.1%, respectively (208, 159, and 86 ASVs). While these phyla generally showed similar broad distribution patterns across each city, *Nitrospirota* were noticeably less abundant in Rome, with median relative abundance approximately two orders of magnitude lower than in the other cities (Figure 1A). At the phylum level, 3.3% of sequences remained unclassified, including two of the top 25 most abundant individual ASVs (ASV20 and ASV17, with 0.72% and 0.7% of all sequences, respectively).

The top fifteen most abundant genera are shown in Figure 1B. Their pattern is more variable, with greater spread among samples within cities. For example, the relative abundance of the top genus *Denitratisoma*, in the *Gammaproteobacteria* class *Burkholderia*, varied widely, from being barely above the limit of detection in some samples, to encompassing over 50% of reads in other samples. Similar differences in relative abundance, several orders of magnitude within the same city, were seen in many of the other top fifteen genera. Although most of these genera were common across the four cities, the *Gammaproteobacteria* genera *Blastomonas* and *Craurococcus-Caldovatus* as well as the *Alphaproteobacteria* genus *Porphyrobacter* were nearly absent from Copenhagen samples but relatively abundant elsewhere.

### Bacterial community diversity

Before looking for specific taxa associated with culturable *Legionella*, we examined how the overall bacterial community structure compared across the four cities, and whether it differed depending on *Legionella* culture status of the samples.

Estimated population richness (number of ASVs) was significantly higher and more variable in Copenhagen and Warsaw, compared with Athens and Rome (Figure 2A, with statistical output in Tables S1 and S2). Taking into account the influence of the city, culture status also had a significant correlation with richness, with culture-positive samples having, on average, 122.0 more estimated ASVs than culture-negative samples (p < 0.001). However, there were significant interactions of city and culture status, indicating that the effect of these factors on richness was not consistent across the four cities. While *Legionella*-positive samples had higher bacterial community richness in Copenhagen, Athens, and, to a lesser extent, in Rome, the opposite was true in Warsaw, where *Legionella*-negative samples had higher richness.

**Figure 2.**
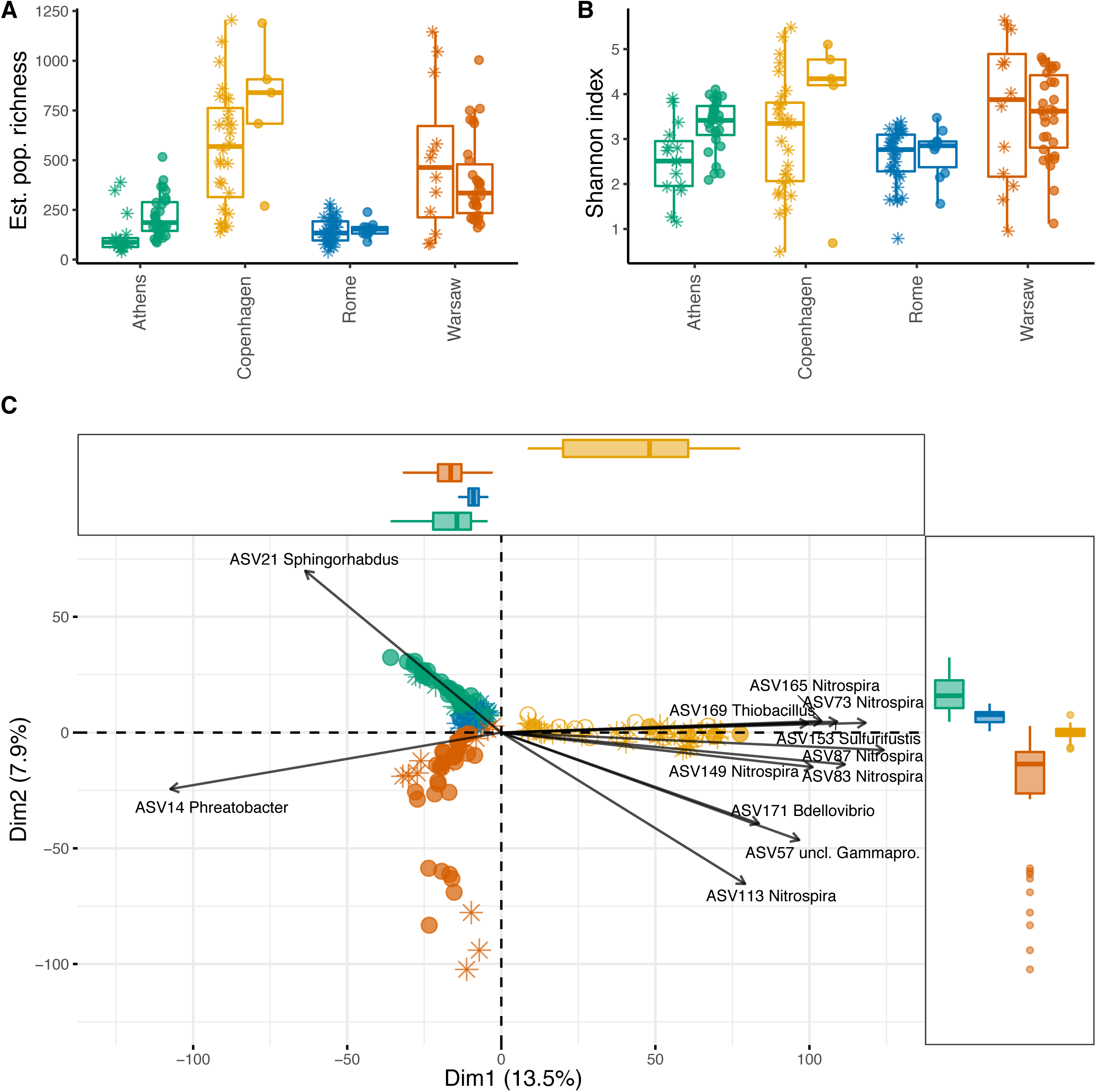
Alpha and beta diversity of premise plumbing samples from four European cities, with and without culturable Legionella. Asterisks show samples that were *Legionella* culture-negative, filled circles show samples that were *Legionella* culture-positive, open circles show samples without *Legionella* culture data. (A) Population richness (number of ASVs) estimated from sample richness using breakaway, which fits nonlinear regression models on the ratios of frequency counts to estimate total richness. This approach does not require sample normalisation, since greater sequencing depth reduces the error of the estimate. (B) Sample Shannon diversity index. Samples were normalised to 10 148 sequences. (C) Aitchison distance PCA, with arrows indicating the fifteen most influential ASVs. Percent variance explained by the first two components is shown on the axes, and marginal boxplots show the distribution of samples along these components.

The Shannon diversity index, a measure of entropy, indicates how evenly the reads are distributed among the ASVs in a sample. Communities with a small number of dominant ASVs would have low Shannon indices, whereas more even community structures would result in higher Shannon indices. Here, despite the significant differences in richness among the cities and between culture-positive and culture-negative samples, Shannon index was similar across samples, indicating a common community structure in these environments (Figure 2B, Tables S3 and S4). Neither city (F3,9.99 = 2.7, p = 0.1), culture status (F1,135.41 = 3.7, p = 0.057), or their interaction (F3,127.92 = 1.0, p = 0.38) had a significant influence on Shannon diversity.

PCA on Aitchison distance clearly clustered the samples by their city (Figure 2C). Copenhagen and Warsaw formed looser, broader clusters than Rome and Athens, indicating more intra-city variability (which likely contributes to the much higher overall richness observed in each of those cities). The percent variance explained by the first two components is shown on the axes. PC1 clearly delineates all Copenhagen samples from the others, suggesting a greater difference in the bacterial communities in those samples compared with the other cities. In contrast, PC2 separates all four cities, pointing to the strong influence of local factors at the ASV level.

The most influential ASVs contributing to the separation of Copenhagen samples along PC1 predominantly belong to the *Nitrospira*, with several ASVs within this genus being highly correlated (i.e. their arrows follow the same path). Copenhagen stood out as having a higher diversity of *Nitrospira* than the other cities, with 676 ASVs falling into this genus, accounting for 2.6% of its 16S rRNA sequences. In comparison, Warsaw, Athens, and Rome had 257, 97, and 33 *Nitrospira* ASVs, respectively, accounting for 1.1%, 1.4%, and 0.08% of sequences from each of those cities.

We used PERMANOVA, as implemented by the adonis function in the vegan package, to test the significance of groupings, blocking by building location ID to account for repeated measures (Table S5). The groupings were significantly influenced by both city (F3,157 = 14.3, R2 = 0.2, p = 0.0014) and *Legionella* culture status (F1,157 = 5.7, R2 = 0.027, p = 0.0005), but the interaction of these variables was not statistically significant (F3,157 = 2.7, R2 = 0.037, p = 0.054).

Since PERMANOVA is sensitive to differences in group dispersions, these were tested using the betadisper function in the R package vegan (Table S6). Group variances were significantly different for city (F3,161 = 15.4, p = 0.0001), indicating that the variability of communities was greater in some cities than in others, which is apparent from the PCA (Figure 2C). The group variances for *Legionella* culture status were not significantly different (F1,163 = 0.077, p = 0.78), so the variability was similar for communities with and without culturable *Legionella*.

Figure 3A shows the intersection of ASVs across the four cities. The rarefied data set used in this analysis (with samples normalised to 10 148 sequences) contained 5086 ASVs. Of these, only 188 (3.7%) were observed in all four cities. Most ASVs, 3545 in total, were detected in only one of the four cities, with 1 827 of these unique to Copenhagen but only 231 ASVs unique to Rome. Not only did the total richness vary markedly between pairs of cities, with Copenhagen and Warsaw having greater richness than Athens and Rome, but the proportion of ASVs unique to each city also differed: 62.9% of ASVs from Copenhagen were not detected anywhere else, whereas these proportions were 47.3% in Warsaw, 34.7% in Athens, and 23.6% in Rome.

**Figure 3.**
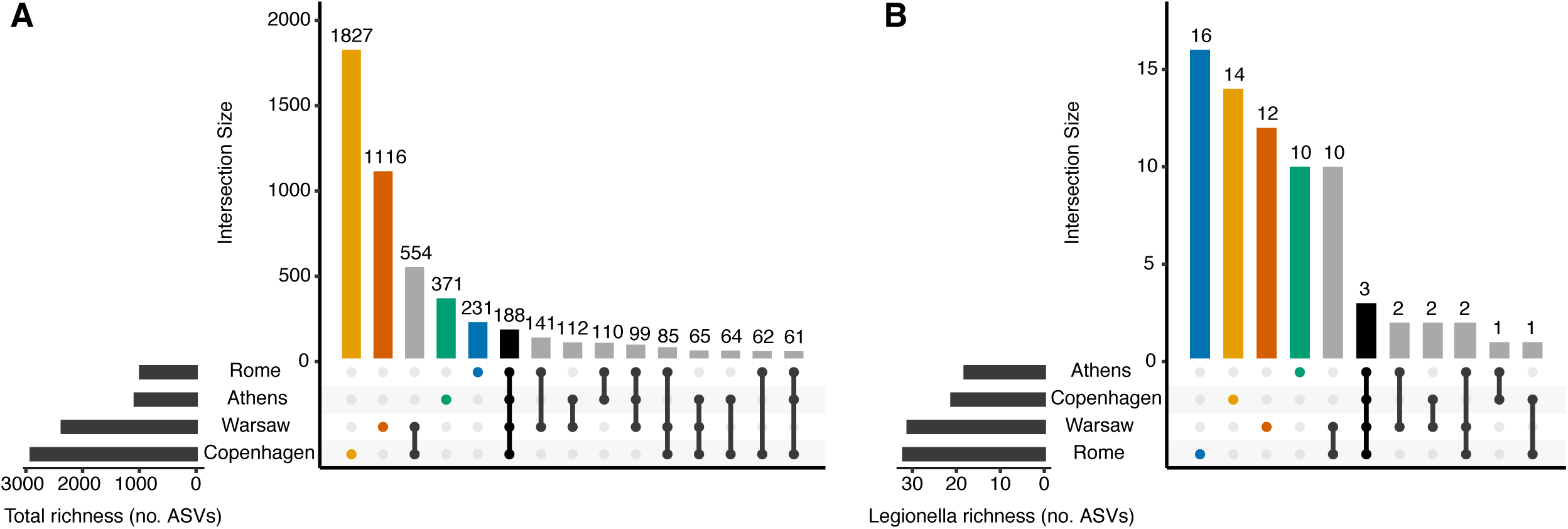
Number of shared and distinct ASVs among the four European cities. Samples were normalised to 10 148 sequences prior to determining the ASVs present in each city, and no minimum prevalence or abundance thresholds were used to determine membership. (A) All 16S rRNA ASVs, and (B) ASVs classified into the genus *Legionella*.

### Legionella ASVs in the 16S rRNA data

In the rarefied 16S rRNA data set, 73 ASVs were classified as *Legionella* at the genus level, accounting for 0.55% of sequences. Seven of these *Legionella* genus ASVs could be further classified to species level. *Legionella* genus ASVs were present in 155 samples overall (88.1%), and in the majority of samples from each city: 77.8%, 95.5%, 91.1% and 88.1% of samples from Athens, Copenhagen, Rome and Warsaw, respectively.

Most of the *Legionella* genus ASVs, 52 out of 73, was unique to one of the four cities. Only three *Legionella* ASVs were detected in all four cities (Figures 3B and 4), and these could all be classified to species level: ASV142 (*L. pneumophila*), ASV425 (*L. anisa*), and ASV1223 (*L. waltersii*). Although Rome had the lowest total bacterial ASV richness and the smallest number of unique bacterial ASVs of any city (Figure 3A), it had the largest richness (32 ASVs) and the greatest number of city-specific *Legionella* ASVs (16) (Figure 3B). Overall, 3.3% of ASVs detected in Rome were classified as *Legionella*, whereas this proportion was 1.7, 1.3, and 0.72% in Athens, Warsaw, and Copenhagen, respectively.

**Figure 4.**
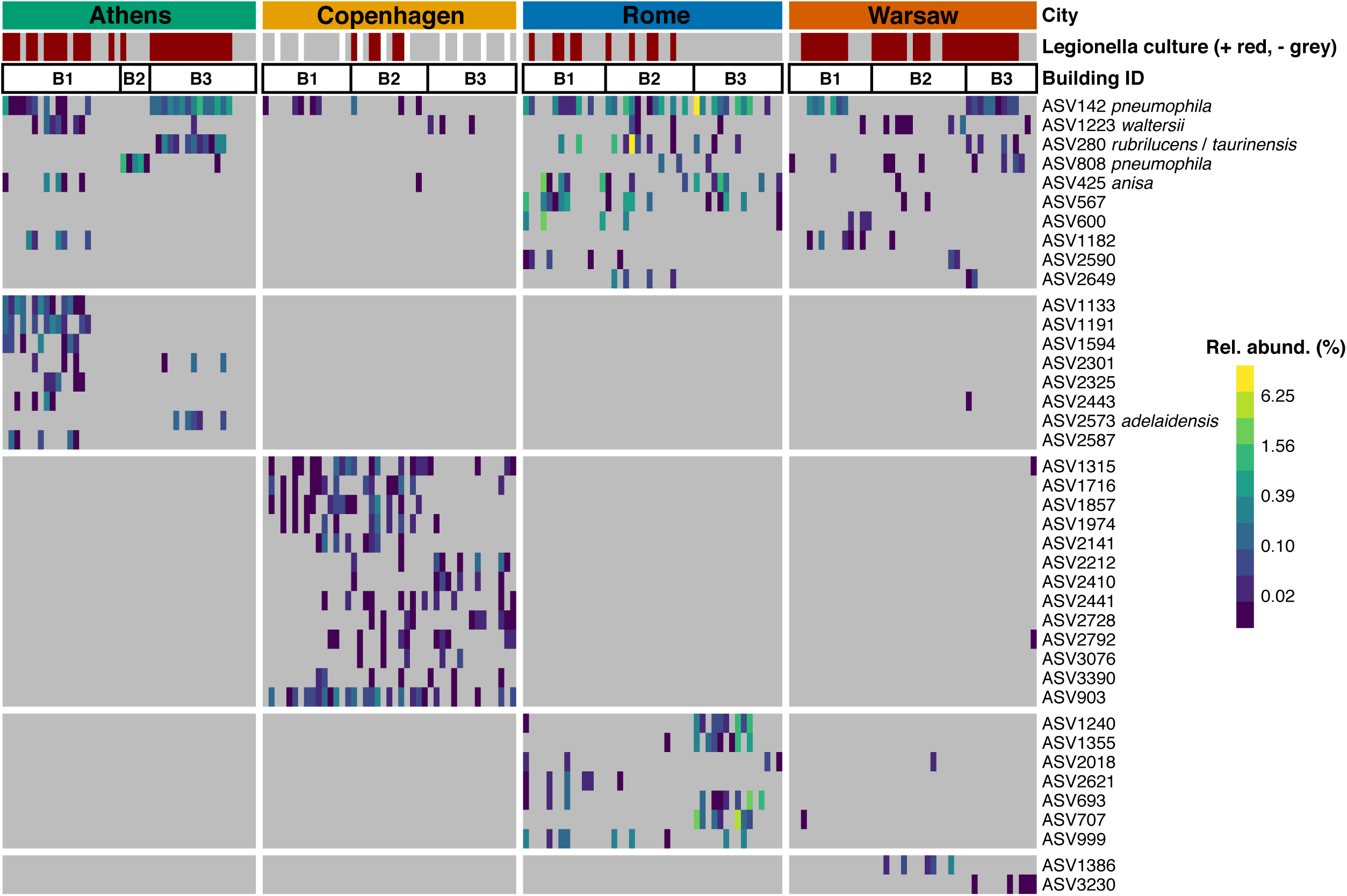
Prevalence and relative abundance of ASVs classifying as Legionella, across all samples, grouped by city and by building. Samples were normalised to 10 148 sequences. *Legionella* culture status of each sample is shown in the bar at the top (red = positive, grey = negative, white = no culture data collected). ASVs are grouped by their prevalence either across multiple cities (top horizontal block), or predominantly in a single city (bottom four horizontal blocks). *Legionella* genus ASVs that could be further classified have species names attached. Only ASVs present in five or more samples are shown, for clarity. Two rarer ASVs were also further classified to species level: *L. drozanskii* (ASV281), present in four samples (two from Rome, two from Warsaw), and *L. pneumophila* (ASV4571), present in one sample from Rome.

The abundance and prevalence of all *Legionella* ASVs present in five or more samples is shown in Figure 4. Most of the *Legionella* ASVs that could be resolved to species level were detected in more than one city, though Copenhagen stood out as rarely having any of these shared, species-resolved ASVs. In many cases, for example in Athens, the specific *Legionella* ASVs detected differed markedly between buildings rather than being distributed evenly across all the samples from that city, and while some buildings (e.g. B1) had a diverse group of *Legionella* ASVs, others (e.g. B2) had only one that was rarely seen elsewhere. A building-specific pattern was less clear in Copenhagen, which had a diverse (albeit low-abundance) population of city-specific ASVs that occurred across all three buildings that were sampled.

Overall, the most abundant *Legionella* genus ASV, ASV142, was further classified as *L. pneumophila*, followed by ASV280, which was unclassified at the species level, though BLAST search against GenBank showed it has 100% similarity to both *L. taurinensis* and *L. rubrilucens*. Two rarer ASVs, ASV808 and ASV4571, were also classified as *L. pneumophila*. The latter was detected in a single sample from Rome, but ASV808, which differs from ASV142 by a single nucleotide, was found in all cities except Copenhagen, though it was particularly prevalent in Athens Building 2, which, unlike all other sampled buildings in this study, had no other *Legionella* sequences (Figure 4).

### Legionella culture counts versus presence in 16S rRNA data

All samples were tested for culturable *Legionella*, except the first 11 samples from Copenhagen, for which *Legionella* culture counts were not collected. Of the 165 samples with culture data, 73 (44.2%) had culturable *Legionella*. The proportion of *Legionella* culture positive samples differed widely among the four cities: 15.2% from Copenhagen, 22.2% from Rome, 62.2% from Athens, and 71.4% from Warsaw. Culture counts in the *Legionella*-positive samples also varied widely, over five orders of magnitude (Figure 5A). The difference in culturable *Legionella* CFU/L was generally not significant among the four cities, except that Copenhagen had significantly lower CFU/L values than Warsaw in pairwise comparisons (zero-inflated GLMM with Poisson distribution, t-ratio153 = -3.023, p = 0.015, Tables S7, S8, S9). The building and compartment random effects improved model fit, as did modelling the zero counts separately by City and random effects, suggesting that the distribution of zero values (i.e. *Legionella* culture-negative samples) differed among the cities (as observed in Figure 5A), and also among the buildings and compartments.

**Figure 5.**
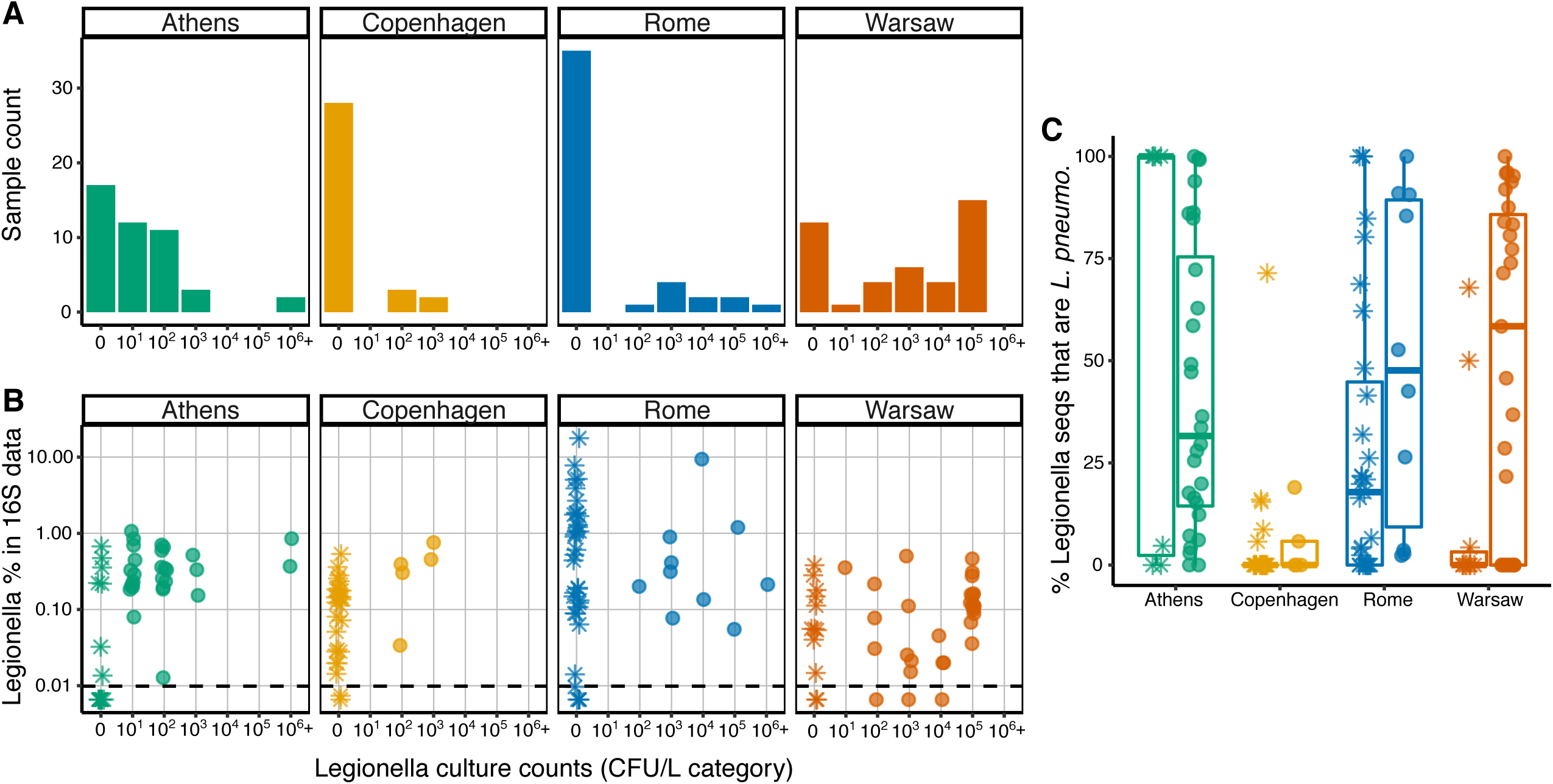
Legionella spp. in the culture and 16S rRNA data. (A) Distribution of samples from each city across the culturable *Legionella* categories (log10 bins, with ‘0’ indicating culture-negative samples. (B) Percent 16S rRNA sequences classifying as the genus *Legionella* versus culturable *Legionella* count categories. Dashed horizontal line shows the limit of detection of the 16S data, 0.0099%, determined from the normalised sequencing depth of 10 148 sequences per sample. (C) Percent *Legionella* spp. 16S rRNA sequences that were further classified as *Legionella pneumophila*. Asterisks indicate samples that were *Legionella* culture-negative, filled circles show samples that were *Legionella* culture-positive.

In three instances, samples were culture-positive but had no detectable *Legionella* spp. in the sequence data (the limit of detection in the sequence data is determined by the sequencing depth, here normalised to 10 148 sequences per sample, giving a LOD of 0.0099%). Conversely, 75 samples were culture negative but had detectable *Legionella* spp. in the sequence data. Seventeen samples were below the detection limits in both culturing and sequencing, whereas 70 had detectable *Legionella* in both approaches.

While the relative abundance of *Legionella* spp. sequences in the 16S rRNA data did not differ significantly across all four cities (F3,9.68 = 3.4, p = 0.063), Rome exhibited a different pattern from the others (Figure 5B). In the other cities, the proportion of *Legionella* sequences in each sample did not exceed 1% (with one exception from Athens at 1.06%), but Rome had 16 samples with *Legionella* above this threshold, including 5 samples with *Legionella* sequences exceeded 5%, with one reaching the highest proportion observed in any sample, 17.7%.

Whether the samples were culture-positive or culture-negative had no significant effect on the proportion of sequences that were placed in the genus *Legionella* (F1,107.8 = 0.1, p = 0.75), and this was the case in all cities, as there was no significant interaction between city and presence/absence of culturable *Legionella* (F1,107.8 = 0.1, p = 0.75). Detailed model outputs are shown in Table S10. The random effects terms did not significantly improve the model (p = 0.51 and 0.49 for building and compartment ID, respectively), indicating that these factors do not explain a significant amount of the variance in *Legionella* sequence proportions. These findings held regardless of whether the culture data were expressed in binary terms (pos/neg), in discrete categorical terms (log10 bins) or on a continuous scale as CFU/L. For subsequent analyses, only the presence or absence of any culturable *Legionella* was considered, not the CFU/L obtained.

Culturing is biased towards *L. pneumophila*, so next, we examined the proportion of sequences that classified specifically as *L. pneumophila*, rather than all those falling into the genus *Legionella*. As a proportion of all 16S rRNA sequences, there was no significant difference in *L. pneumophila* among the cities or between culture-positive and culture-negative samples (Table S10).

However, differences were observed when the *L. pneumophila* proportion was calculated not from all 16S rRNA sequences, but from those classifying as the genus *Legionella* (Figure 5C). Copenhagen had the lowest proportion of *L. pneumophila*, indicating that the majority of *Legionella* sequences detected in that city were from other species (as also seen in Figure 4). In the other cities, some samples had only *L. pneumophila* sequences, some had no *L. pneumophila* sequences among their *Legionella* spp. data, and many had a mix of *L. pneumophila* and other species. Given the variability, the differences among the cities was not significant (F3,8.03 = 1.9, p = 0.21), but taking into account the influence of the city and the random effects (building and compartment), the culture status of the sample (*Legionella*-positive or negative) had a significant influence on the proportion of *Legionella* genus sequences that were further classified as *L. pneumophila* (Table S11), with the proportion in culture-positive samples being, on average, 24.7% higher than in culture-negative samples.

Despite this agreement between culture status and proportion of *Legionella* sequences classified as *L. pneumophila*, we used the culture status rather than sequence data to delineate between samples with and without *Legionella* to address our main question: whether other members of the bacterial communities are associated (positively or negatively) with *Legionella*. Sequencing total environmental DNA can detect dead, nonviable, and dormant cells, whereas culturing confirms the presence of viable *Legionella*.

### Associations of bacterial taxa with Legionella

Despite the variability in the sample types and the differences in community diversity among the four cities, differential abundance analysis was able to identify 54 specific ASVs (Figure 6A) that are negatively or positively associated with culturable *Legionella*. Only ASVs present across a relatively large number of samples could show significant associations, and given the variability among cities at the ASV level, a relatively small number met this conditions. We therefore also clustered into genera to identify taxa that might fulfill similar roles in different places, despite regional variations in the exact species and strains within those genera. We identified 42 specific genera (Figure 6B) with a significant association, not including those unclassified at the genus level (since the agglomeration step risks grouping together quite different unclassified taxa that share the family level).

**Figure 6.**
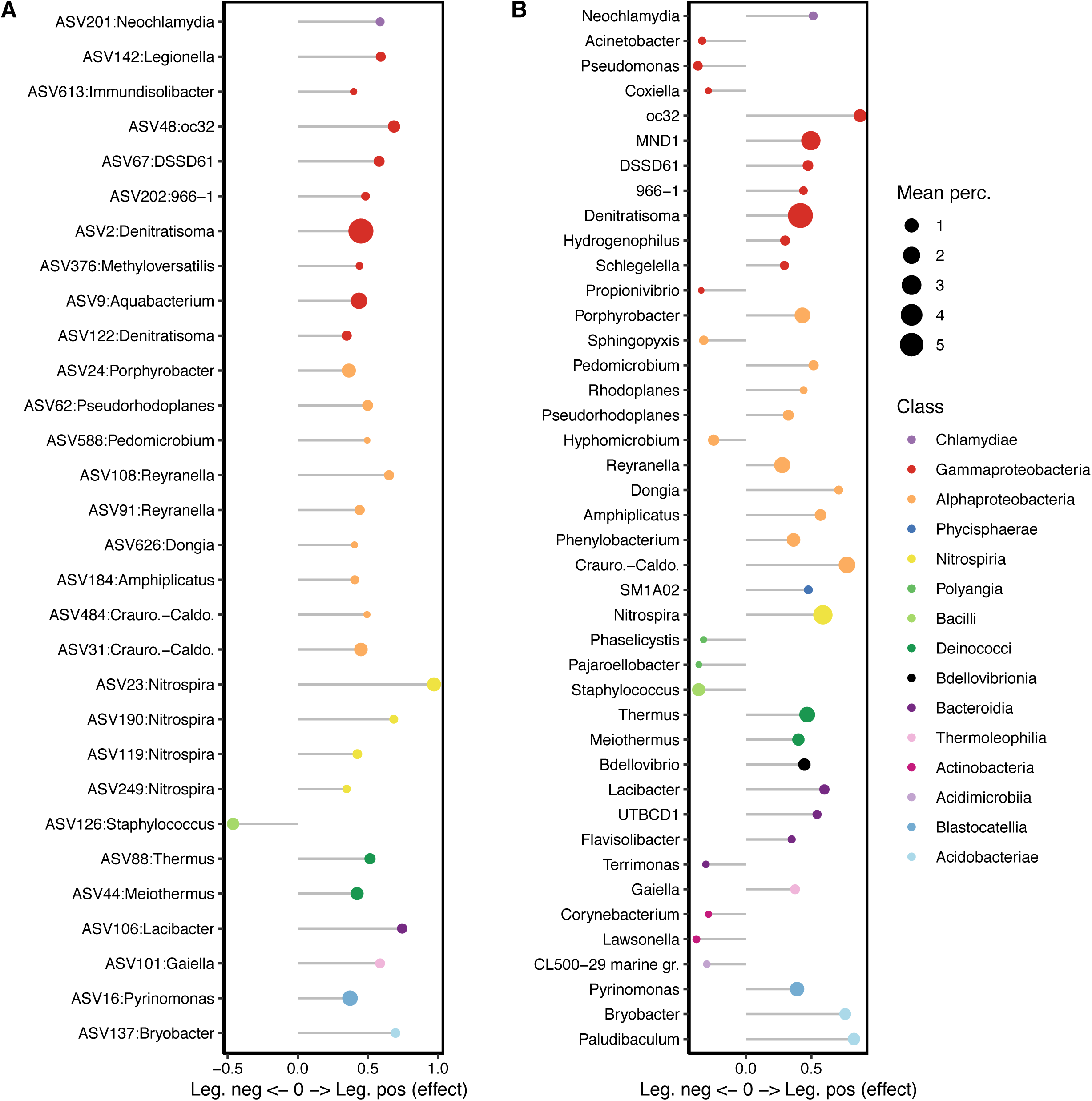
Specific ASVs (A) and genera (B) showing a significant association with Legionella culture-negative or culture-positive samples. ASVs and genera with effect < 0 are significantly associated with culture-negative samples, and those with effect > 0, with culture-positive samples. Symbols are coloured by Class and scaled to the mean relative abundance of the ASV or genus across all samples.

### Negative associations

At the ASV level, only ASV126 was identified as having a significantly negative association with culturable *Legionella*. It was classified as genus *Staphylococcus* (Phylum *Firmicutes*, Class *Bacilli*), but could not be resolved to species level as it showed 100% identity to *S. capitis*, *S. epidermidis*, and *S. capra* when searched against GenBank.

When ASVs were grouped at the genus level, more negative associations were identified, with 13 named genera having a significant negative association with culturable *Legionella* (Figure 6B). These include the *Gammaproteobacteria* genera *Pseudomonas* and *Acinetobacter* (Figure 7A), the *Alphaproteobacteria* genera *Hyphomicrobium* and *Sphingopyxis*, as well as the genus *Staphylococcus*.

**Figure 7.**
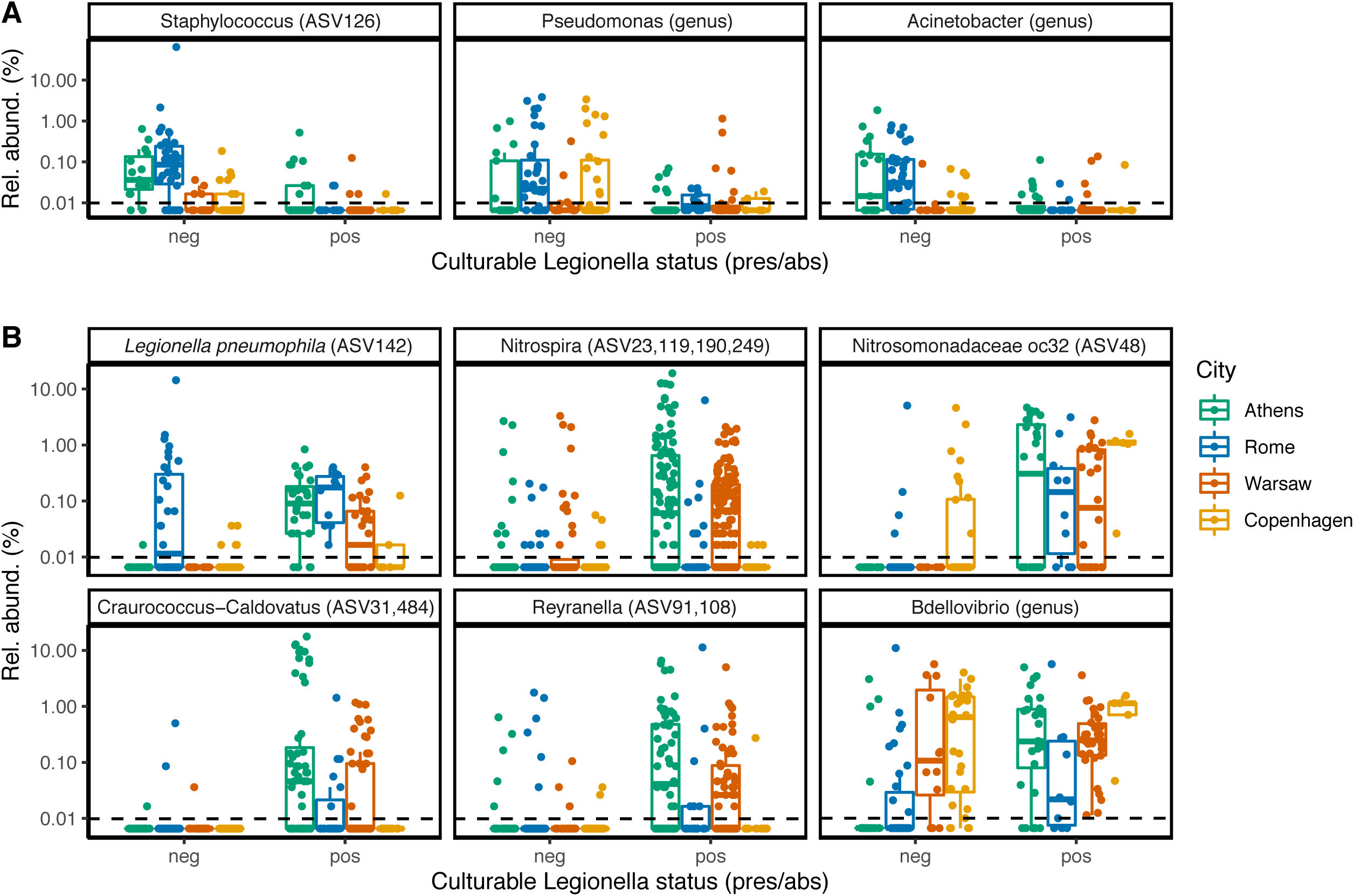
Examples of differentially abundant ASVs and genera. (A) Taxa associated with *Legionella* culture-negative samples. (B) Taxa associated with *Legionella* culture-positive samples. Differential abundance analysis was carried out separately on the ASV-level and genus-level data, using ALDEx2.

### Positive associations

Numerous individual ASVs and genera were positively associated with culturable *Legionella* (Figure 6). At the ASV level, ASV142, *Legionella pneumophila*, had a positive association overall despite its unusual abundance in the culture-negative samples from Rome. Even though the proportion of all *Legionella* genus sequences could not be tied to culturable status, this specific *L. pneumophila* ASV could.

Of the 59 ASVs classified as the nitrite oxidising genus *Nitrospira*, four were positively associated with culturable *Legionella*, and their combined relative abundance across the cities is shown in Figure 7B. Despite the genus *Nitrospira* forming a large proportion of the bacterial communities in Copenhagen, more so than in the other cities, these four specific *Legionella*-associated *Nitrospira* ASVs were hardly detected in samples from there (Figure 7, Figure S3), and they were not among the *Nitrospira* ASVs that contributed to the separation of Copenhagen samples in the PCA (Figure 2C). The specific *Nitrospira* ASVs that were abundant in Copenhagen were not positively associated with *Legionella*.

At the genus level, 29 genera had a positive association with culturable *Legionella*. These included the genus *Nitrospira* overall, as well as the nitrifying genus *oc32*, from the family *Nitrosomonadaceae*. *Craurococcus* and *Reyranella*, genera known to be associated with amoeba, were also significantly associated with *Legionella* culture-positive samples, as were *Bdellovibrio* and the *Acidobacteria* genera *Paludibaculum* and *Bryobacter*.

## Discussion

*Legionella* proliferation in engineered water systems, including premise plumbing, is a wide-scale problem with measurable impacts on human health. Understanding the ecology of *Legionella* in these systems, notably its interactions with other microbial community members, is key to developing more robust and sustainable control measures. Here, we showed that numerous bacterial community members displayed significantly different abundance patterns depending on whether culturable *Legionella* were present or not in a premise plumbing water sample. Taxa that showed a positive association with culturable *Legionella* included members of ameoba-associated genera (*Reyranella* and *Craurococcus-Caldovatus*), those involved in nitrogen cycling (*Nitrospira*, *Denitratisoma*, *Nitrosomonadaceae*), and the predatory genus *Bdellovibrio*. Other taxa, notably *Staphylococcus*, *Pseudomonas*, and *Acinetobacter*, were more abundant in samples without culturable *Legionella*, suggesting a possible inhibitory effect. These positive and negative relationships held across the four European cities and across the different buildings and plumbing compartments. This is important because developing better control measures, including probiotic approaches, will require an understanding of ecological relationships that can be generalized across different systems.

The differences between culture-positive and culture-negative samples were not limited to individual bacterial genera and ASVs. At the community structure level, although we found no significant differences in Shannon index between culture positive and negative samples, pointing to similar dominance/evenness structures (also reported in Canadian cooling towers (17)), the estimated community richness differed significantly, with higher richness overall in *Legionella* culture-positive samples. The same observation was reported in cooling towers across the USA, where bacterial community richness was positively correlated with detectable *Legionella* (19). The USA premise plumbing study limited its analysis to the Class level, so specific families or genera associated with *Legionella* could not be resolved, but both cooling tower studies showed that *Pseudomonas* (or the family level, *Pseudomonadaceae*, in the USA study) had a negative correlation to *Legionella* spp., suggesting that the inhibitory effect seen in laboratory studies might also be occurring in cooling towers. The only other taxon that showed a negative relationship specifically to Lp, reported in the Quebec study, was the genus *Sphingobium*. The same study reported that numerous genera were enriched in *Legionella*-positive samples, including *Porphyrobacter*, *Sediminibacterium*, *Sphingopyxis*, *Hyphomicrobium*, and *Reyranella*, among others (17).

Why greater community richness is associated with a higher probability of growing *Legionella* is unclear. It could point to a lower concentration of residual disinfectant in those compartments, allowing not only *Legionella* but other taxa to flourish. This would not, however, explain the difference observed in Copenhagen, since no disinfection is used there but *Legionella*-positive samples had, on the whole, richer communities than *Legionella*-negative ones. Perhaps the greater community richness in culture-positive samples is due to *Legionella*’s requirement for specific amino acids, which might be in greater supply in richer communities, where several different taxa can supply them. However, the effect of *Legionella* culture status on richness in our study was complicated by a significant interaction with the city, meaning the effect was not observed equally across all cities. Indeed, it was reversed in Warsaw, where culture-negative samples had higher estimated richness.

The differential abundance signal from positively and negatively associated taxa was detectable despite there being significant differences in premise plumbing bacterial communities across the four cities, with relatively little overlap in ASVs, most of which were city-specific. Total richness and number of city-specific ASVs were highest in Copenhagen and Warsaw, but much lower in Athens and Rome. Drinking water systems without a disinfectant residual are known to be richer than those with a residual (21, 22), so the absence of chemical disinfection in Copenhagen’s public water supply likely explains why it had the richest premise plumbing communities, the largest number of city-specific ASVs, the greatest dispersion among samples, and the greatest beta diversity separation from the other cities, accounting for 13.5% of variance across the data set. A possible reason of such high richness of Warsaw compared with Athens and Rome, although the water is treated with chlorine dioxide, it is supplied originally from the Vistula river and the Dębe Dam lake that may be microbiome rich bodies of water.

The coarse taxonomic composition of premise plumbing communities was similar in these four cities, with strong dominance of *Proteobacteria* classes *Gamma-* and *Alphaproteobacteria*. This is in line with other studies of water distribution systems (22). At finer taxonomic resolution, there were some noticeable differences among the four cities, with *Nitrospira* abundance and diversity being much higher in Copenhagen than elsewhere, but *Craurococcus-Caldovatus* and *Blastomonas* being almost absent from there despite being relatively abundant in the other cities. Furthermore, the abundance and diversity of sequences classifying as *Legionella* varied across the four cities. Rome had the highest number of *Legionella* ASVs overall, the highest number of city-specific *Legionella* ASVs, and the highest proportions of *Legionella* in the 16S rRNA sequence data, with many samples exceeding 1% (with one sample having 17% of sequences classified as *Legionella*). Additionally, many of the relatively abundant *Legionella* ASVs in Rome were named taxa, including Lp, which are more likely to be involved in human disease than rare, uncharacterised species. In contrast, most *Legionella* ASVs detected in Copenhagen were at low relative abundance, were unnamed and uncharacterised, and were city-specific. There was a near absence of the main named species that cause most human illness. The *Legionella* ASV findings in these two cities sit in contrast to the total community richness and diversity, which was low in Rome and high in Copenhagen. This suggests that rich bacterial communities provide some competition against or dilution of *Legionella* species, but this would contradict our finding that samples with richer total communities were more likely to have culturable *Legionella* (albeit not in all cities).

This contradiction could be due to the absence of a correlation between culturable *Legionella* and abundance in the 16S rRNA sequence data. A similar mismatch between *Legionella* as detected by PCR versus in pure culture was also observed in US cooling towers (19). In our study, this mismatch was especially evident in Rome, which had a relatively high abundance of *Legionella* 16S rRNA sequences, even in samples that failed to grow *Legionella* (which included the sample with 17% *Legionella* sequences). Why such a mismatch was observed is unclear; it could be due to disinfection regimes that kill cells after a period of proliferation, without destroying their DNA, or that cause a switch to a viable but nonculturable state, meaning they would be detected by molecular but not culture methods. Culturing is the gold standard, and detection of *Legionella* DNA is not necessarily a good indicator, since it cannot distinguish between loss of viability and viable but nonculturable state (19). For our differential abundance analyses, culturable *Legionella* status, not presence in sequence data, was therefore used to identify negative and positive associations of other community members.

Several of the negative associations that we identified are well supported by laboratory studies. For example, we found that a specific *Staphylococcus* sequence (ASV126) as well as the genus *Staphylococcus* overall were enriched in *Legionella* culture-negative samples, and pure culture experiments have shown that members of this genus can produce anti-*Legionella* peptides (8, 23). However, this negative association was not reported in the cooling tower studies, as *Staphylococcus* species were not detected there (17, 19). The negative effect of *Pseudomonas* on *Legionella* proliferation is well documented from laboratory studies, with *Pseudomonas* species using numerous inhibitory mechanisms (9–13). The cooling tower association studies in Canada (17) and across the USA (19) also found that the genus *Pseudomonas* and the family *Pseudomonadaceae*, respectively, had a negative correlation with *Legionella*, and we detected the same pattern on another continent and in premise plumbing compartments, which are different types of engineered water systems. This suggests that the inhibitory effects of *Pseudomonas* species on *Legionella* transcend the laboratory and can be generalised across multiple types of engineered water systems. The negative relationship in cooling towers was thought to be due (at least in part) to chlorination, which kills *Legionella* but allows *Pseudomonas* to proliferate (17). However, this explanation does not entirely fit our data, because in Copenhagen, there is no chlorination but the observation still holds (there were more *Pseudomonas* in culture-negative samples).

While the *Staphylococcus* and *Pseudomonas* negative associations were supported by published laboratory and/or cooling tower association studies, other negative relationships identified in our data were the opposite of what was reported in cooling towers. *Sphingopyxis* and *Hyphomicrobium* (both *Alphaproteobacteria*) were positively associated with *Legionella* in cooling towers (17) but negatively associated in our data. These genera are particularly abundant in biofilms and are thought to help in biofilm formation (24, 25), so an interesting negative association which may relate to the more organically rich environments of cooling towers and may be less pertinent in nutrient poorer premise plumbing systems that are continuously flushed. In addition, based on laboratory experiments, *Bdellovibrio* should have had a negative influence on *Legionella* proliferation, but was instead positively associated with culturable *Legionella*. *Bdellovibrio* is a predatory taxon but is not a specific grazer of *Legionella* (6), so perhaps it was instead grazing on *Legionella*’s competitors in the biofilm. This highlights that real-world effects are more complex than laboratory experiments. Seeding systems with *Bdellovibrio* to control *Legionella*, for example, could have unintended consequences if complex ecological interactions are poorly understood.

The positive association of *Bdellovibrio* was the opposite of what laboratory experiments would have suggested, but other positive associations were easier to explain. Several were taxa known to be associated with amoebae, notably species in the *Craurococcus-Caldovatus* genus grouping (26) and in *Reyranella*. The positive association of these genera with culturable *Legionella* likely points to a higher abundance of amoebae in general, and thus a greater availability of host cells for *Legionella* growth and replication. A positive association of *Reyranella* was also reported in the Canadian cooling tower study (17). Another amoeba-associated taxon, the symbiont *Neochlamydia*, also showed a positive association with *Legionella*; however, this correlation is more puzzling, as *Neochlamydia* species can confer resistance of their host amoebal cell to infection by *Legionella* and would be expected to have a negative influence (6). In our data, this positive association might simply be highlighting an abundance of amoeba in general.

Several other taxa that were positively associated with culturable *Legionella* appear to be important in nitrogen cycling, which was not apparent in the cooling tower association studies (17, 19). A number of ASVs in the nitrite-oxidising genus *Nitrospira*, as well as the genus overall, showed a strong positive association. This was despite the abundance and diversity of *Nitrospira* ASVs in Copenhagen samples, which only rarely grew *Legionella*. The specific *Nitrospira* ASVs that were most influential in separating the Copenhagen samples from the other cities in the PCA were not the same *Nitrospira* ASVs that had a positive association with *Legionella*. This suggests that even within the same genus, taxa (species, subspecies) can have different types of community interactions, with some species exhibiting a positive association with *Legionella* and others not. In the case of *Nitrospira* at least, but perhaps more generally, the genus level may be too coarse for differential abundance analysis and could mask contrasting patterns at finer taxonomic levels.

Of course, association studies like ours show only correlation, not causation. The taxa with significant differential abundances are not necessarily supporting the proliferation or inhibiting the growth of *Legionella* directly. These taxa, including *Legionella*, could all be responding to other local conditions, for example temperature or disinfectant regime. The scale must also be considered in studies like this; relationships that are statistically significant across cities do not necessarily hold in individual biofilms. Finally, while culture data are quantitative (counts per L), 16S rRNA sequence data can only be semi-quantitative, as they are compositional in nature. They indicate the proportion of the bacterial community accounted for by each ASV, but do not give any indication of the total microbial biomass in these samples, which would have had to have been measured in parallel using a different method (e.g. flow cytometry, qPCR), or estimated using a spike-in control, neither of which we included here but which would be useful to consider in future studies of this nature.

## Conclusions

We found that significant associations (negative and positive) of specific bacterial taxa with culturable *Legionella* could be identified across a highly variable set of samples covering different premise plumbing compartments, different buildings, in four cities across Europe. Some of these associations were previously reported in cooling towers in North America and are supported by laboratory experiments with pure cultures. Our study suggests some of these associations are therefore consistent across many different environments where *Legionella* is found. Identifying ecological associations that transcend the laboratory and apply to different engineered water systems is important for developing new approaches to *Legionella* control.

## Materials and Methods

### Sampling locations and sample collection

Four European capital cities were included in this study: Athens (Greece), Copenhagen (Denmark), Rome (Italy), and Warsaw (Poland). In 2015, Italy and Denmark had population LD incidence rates of >2 cases per 100 000, whereas the incidence rates in Greece and Poland were 0.01-0.49 cases per 100 000 (27). The four cities differ in their water treatment approaches, the most notable difference being that Copenhagen, unlike the other three cities, does not use chlorination in its drinking water distribution system.

Three buildings were selected from each city; this included a mix of hospitals, hotels, healthcare facilities, and public buildings, some of which had additional water treatment systems in place. Within each building, four premise plumbing compartments were chosen for repeated sampling: municipal water at the entrance of each building (MW), boiler bottom (BB) or after heater exchanger (AHE), hot recirculating water (HRW), and the farthest point from the boiler or heat exchanger (FPB). Repeated samples of each compartment were collected four (Athens, Copenhagen, Rome) or five (Warsaw) times between winter 2016-2017 and spring 2018.

Two litres of water were collected from each compartment at each sampling point. One litre was concentrated to 10 mL using sterile 0.22 μm polycarbonate filter membrane for *Legionella* enumeration by culture, following the ISO 11731:2017 protocol. The second litre was filtered through a sterile 0.22 μm polycarbonate membrane, which was used for DNA extraction. Water temperature and pH were measured in a subset of samples.

### DNA extraction and 16S rRNA amplicon sequencing

Genomic DNA was extracted using the PowerWater DNA isolation kit (Qiagen, Hilden, Germany), according to the manufacturer’s instructions. Briefly, the filter membrane was inserted into a bead beating tube containing lysis buffer. After disruption of the cells, lysates were transferred to a DNA-retaining spin column for the purification steps, performed in a QIAcube instrument. DNA was eluted in 100 μl elution buffer (10 mM Tris pH 8.5).

High-throughput sequencing of the bacterial 16S rRNA gene V3-V4 hypervariable region was performed by a commercial genomics service (Eurofins Genomics, https://www.eurofinsgenomics.eu). V3-V4 amplicons were generated using the Eurofins standard primers V3V4-F (TACGGGAGGCAGCAG) and V3V4-R (CCAGGGTATCTAATCC). Libraries were cleaned, quantified and pooled for sequencing on the Illumina MiSeq platform with v3 chemistry (2x300 bp).

### Bioinformatics processing

Demultiplexed fastq files obtained from Eurofins were first grouped by the instrument and run number specified in the sequence identifier lines, to account for the different sequencing error profiles that can occur on separate instruments and runs. DADA2 v1.16.0 (28) was used for quality control, error correction, taxonomic assignment, and generation of an amplicon sequence variant (ASV) table. Briefly, sequences were truncated based on their quality profiles, with settings specific to each sequencing run, and filtered to remove those with ambiguous and low-quality bases (full details in the R scripts available on GitHub). After the error modelling step, sample inference was run with the pool=pseudo option to maximise detection of rare variants while remaining within the limits of our computing resources. Paired reads were then merged, discarding any that did not overlap by at least 25 bases with no mismatches. With this primer set, merged reads show a bimodal length distribution with modes at 403 and 427/428 base pairs. Sequences outside the size range 400-430 bp were removed. ASV tables from the different sequencing runs were merged, chimeras were identified and removed using the “consensus” method in DADA2, and taxonomy was assigned using DADA2’s implementation of the native Bayesian classifier (29), against the Silva NR database, v1.38 (30, 31). Further classification to species level was attempted using the Silva v1.38 species training set.

The R package phyloseq (32) was used for further pruning and filtering to remove likely artefacts and non-target sequences. Sequences classifying as Archaea, Eukarya, or unclassified at the Domain level were removed, as were those identified as mitochondria or chloroplasts. A prevalence filter was applied to keep only ASVs observed in at least three samples, and samples with fewer than 2000 reads were discarded.

### Statistical analysis

For most analyses, we used compositional methods that do not require normalisation of sequencing depth (33). However, where normalisation was required, e.g. sample Shannon index, comparison of ASV intersections, and proportions of taxonomic groups, we subsampled (with replacement) down to 10 148 sequences per sample, discarding the samples smaller than this in order to maximise the amount of data retained.

Linear and generalised linear mixed effects models (R packages lme4 (34), lmerTest (35), and glmmTMB (36)) were used to account for the hierarchical and repeated measures structure of the data, and for the zero-inflated Poisson distribution observed with the *Legionella* culture count data. City, and where relevant, presence/absence of culturable *Legionella*, were specified as fixed effects. The repeated measurements within each compartment were accounted for by modelling building compartment as a random effect, nested within the building ID, since compartments within a building might have more similar microbiomes due to a common water source and local disinfection regime. The function ranova in the lmerTest package was used to compute ANOVA-like tables to assess whether these random effect terms improved model fit, and pairwise comparisons among the four cities were computed from the model output with the lsmeans package (37).

For alpha diversity analyses, population richness (the number of ASVs in the populations from which samples were taken, rather than the number of ASVs in the samples themselves) was estimated using the R package breakaway (38), and hypothesis testing was carried out with the functions betta and betta_random (39), which account for the uncertainties in the richness estimates. Fixed and random effects were as above. Sample Shannon index (at the ASV level) was calculated in phyloseq. The intersection of ASVs among the four cities was plotted using the R package UpSetR (40).

For beta diversity analyses, we used the compositional Aitchison distance (33), equal to the Euclidean distance calculated from a center-log-ratio (CLR) transformed ASV table. Prior to the CLR transform, which cannot be done on zero values, we applied the count zero multiplicative approach implemented in the zCompositions package (41). Samples and taxa were ordinated by PCA and plotted using the factoextra package (42). The influence of city and presence/absence of culturable *Legionella* (CFU_pres_abs) was assessed by permutational multivariate analysis of variance (PERMANOVA), as implemented by the adonis function of the vegan package (43). Compartment ID was specified as a blocking variable (strata) to account for repeated measures. Multivariate homogeneity of group dispersions was tested for city and *Legionella* pres/abs with the betadisper function.

Differential abundance analysis was carried out on the ASV and genus-agglomerated tables using the compositional approach implemented in ALDEx2 (44, 45). Presence/absence of culturable *Legionella* was specified as the fixed effect, and significance testing used a Wilcoxon Rank Sum test and Welch’s t-test, with Benjamini-Hochberg corrected p-values.

### Data availability

Raw sequencing data were deposited in the European Nucleotide Archive (ENA) under BioProject PRJEB52062.

## Acknowledgements

This work was funded by ESCMID as the first grant (no number is available) provided to ESGLI (ESCMID Study group of Legionella Infection) in 2016. We wish to thank Dr. Rodrigo Bacigalupe from University of Edinburgh, for his critical evaluation of the paper. We are grateful to Mrs Despina Karamousa from the Region of Attica, Regional Unit of the Central Athens Sector, Department of Health and Environmental Control and Mrs Aspasia Gatsiou from the Specialized Oncologic Hospital of Piraeus “METAXA,” for their valuable collaboration, continuous support and commitment to the project and the laboratory. Finally, we are grateful to the many individuals who have taken the time to publish open-access tutorials, bioinformatics workflows, scripts, R packages, and other resources for the benefit of the scientific community.

---

**Figure S1.**
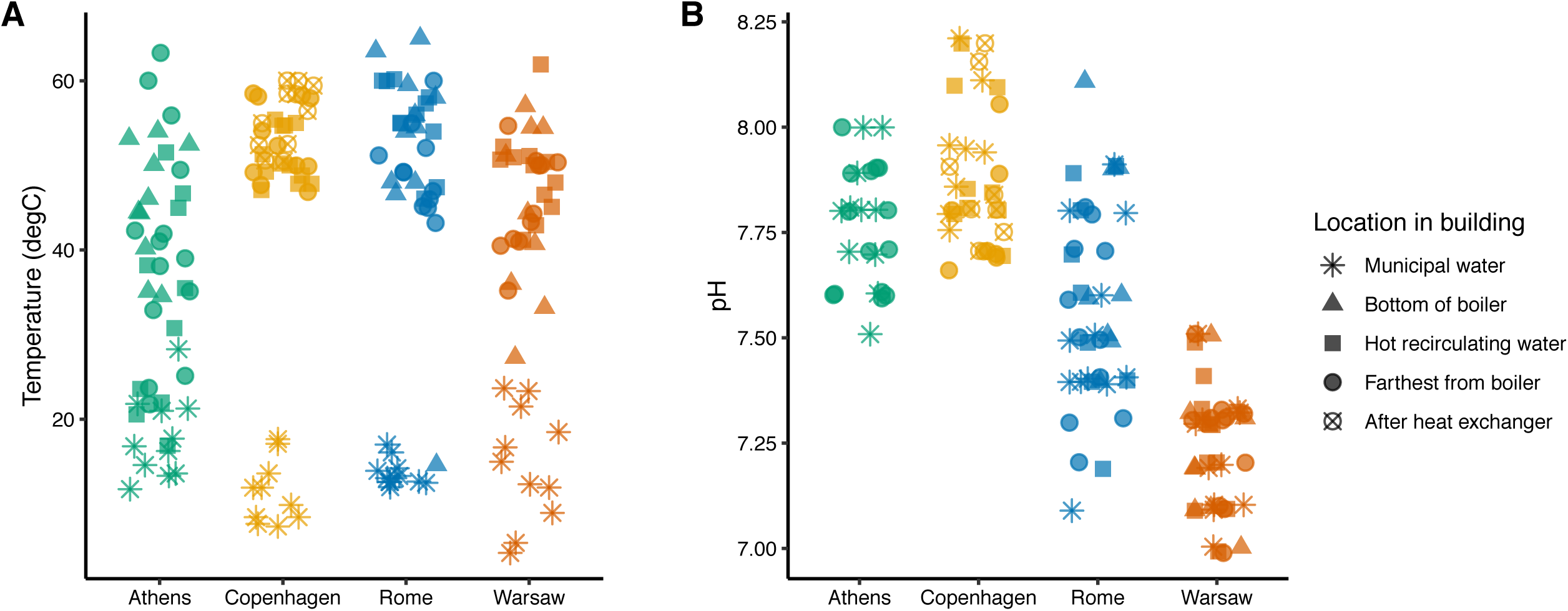
Temperature (A) and pH (B) of water samples collected from different premise plumbing compartments, in three buildings from each of four European cities.

**Figure S2.**
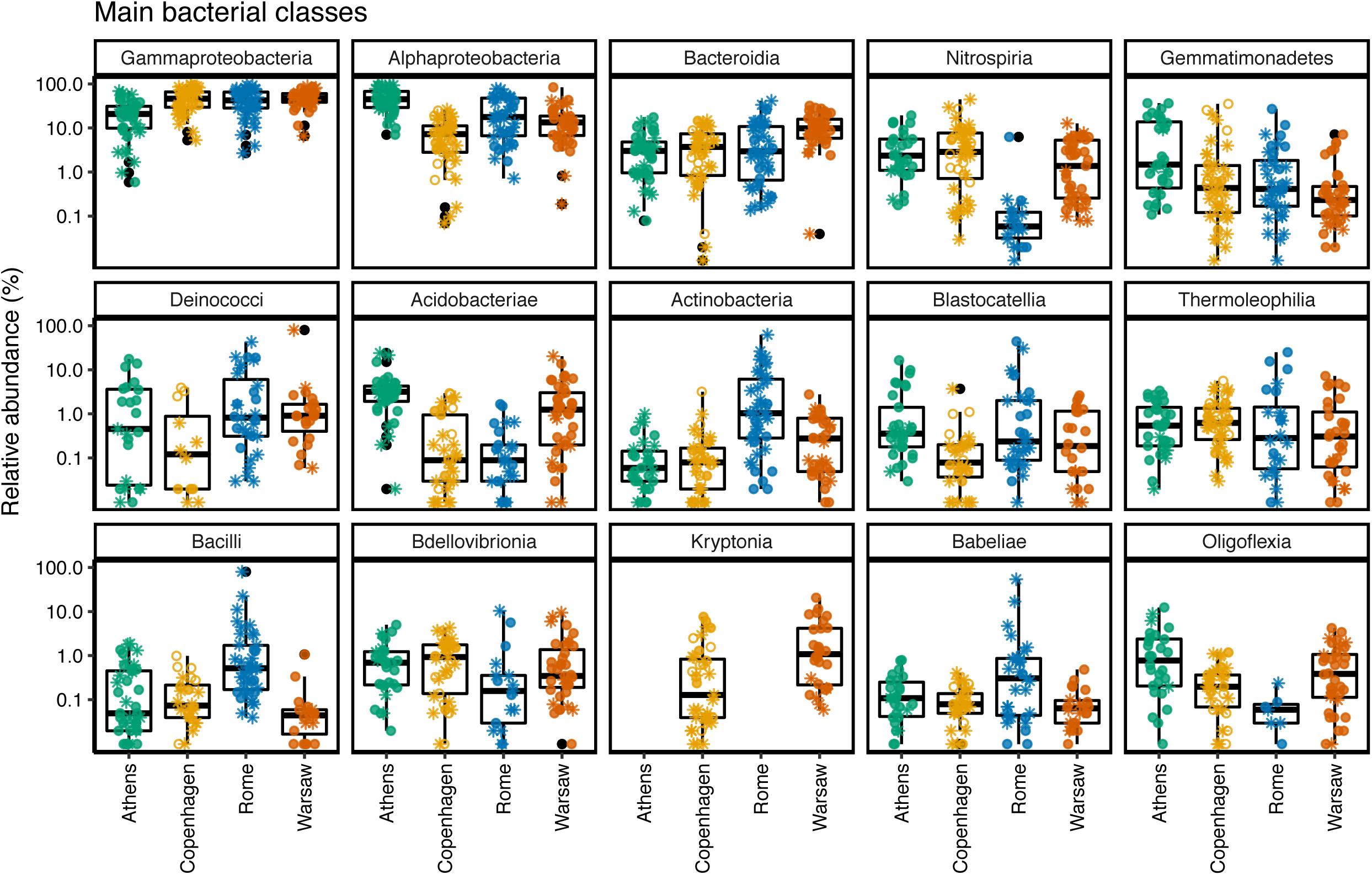
Top fifteen most abundant classes across all samples. Samples were normalised to 10 148 sequences. Asterisks indicate samples that were Legionella culture-negative, filled circles show samples that were Legionella culture-positive, open circles show samples without culture data.

**Figure S3.**
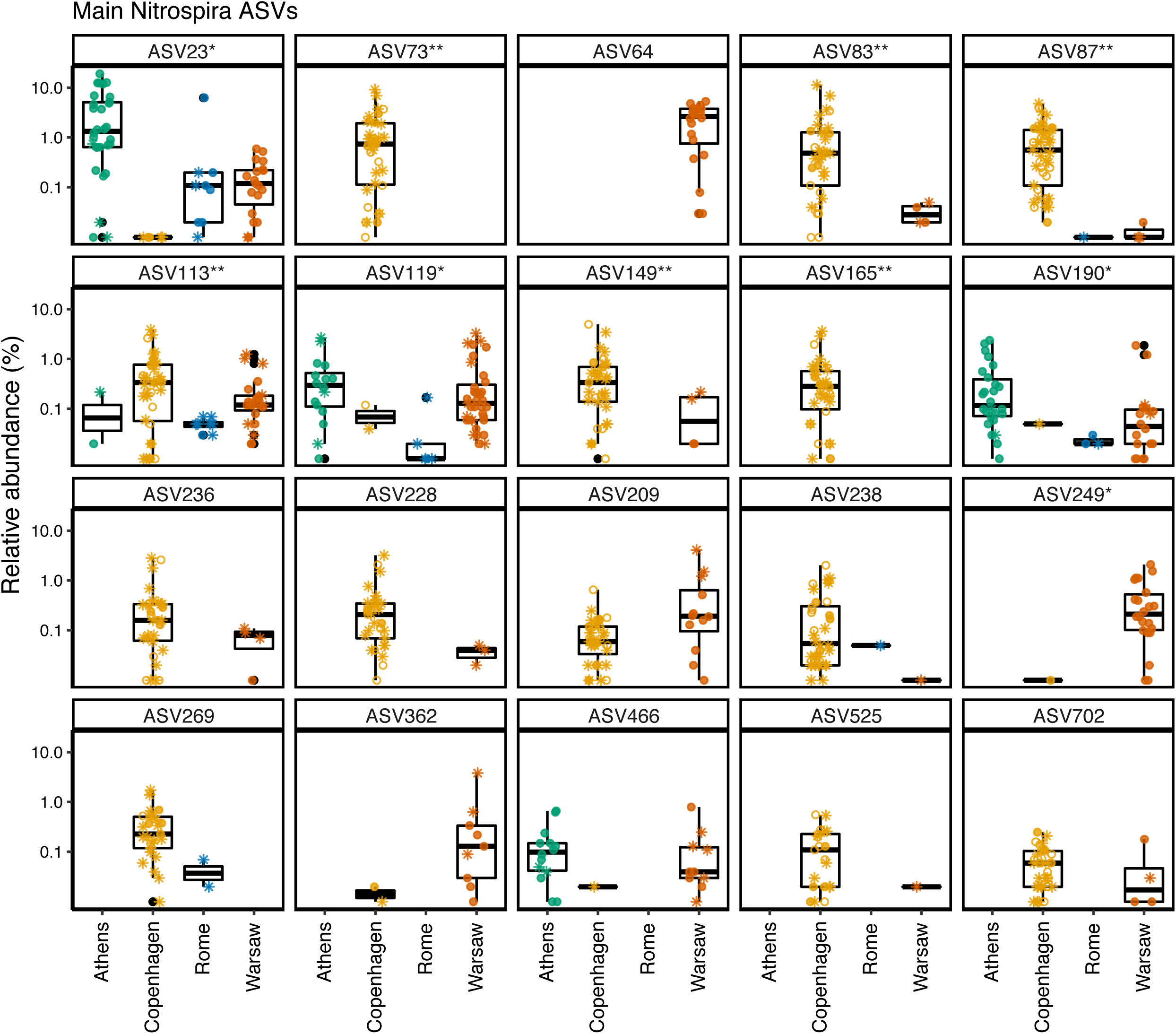
Top twenty ASVs classified as the genus Nitrospira. Samples were normalised to 10 148 sequences. Asterisks indicate samples that were Legionella culture-negative, filled circles show samples that were Legionella culture-positive, open circles show samples without culture data. *ASVs that had a positive association with culturable Legionella, **ASVs among the top fifteen most influential in PCA showing separation of Copenhagen samples from the others.

## Supplemental data

### Statistical tables

#### ASV population richness estimates

Estimated population richness was calculated using breakaway (version 4.7.5). Mixed effects model was run with betta_random, which takes into account the uncertainty around the population richness estimate. City and presence/absence of culturable *Legionella* were specified as fixed effects (including the interaction), and building compartment ID was specified as a random effect (betta_random accepts a single random effect term).

**Table S1.** Population richness: Output of mixed effect model run with the breakaway package, which accounts for the uncertainty around the richness estimates. Model formula: Estimated.richness ∼ City * CFU_pres_abs + (1 | Bldg_loc_ID)

|  | Estimates | Standard Errors | p-values |
| --- | --- | --- | --- |
| (Intercept) | 97.498 | 11.997 | 0.000 |
| CityCopenhagen | 449.793 | 42.239 | 0.000 |
| CityRome | 43.083 | 20.113 | 0.032 |
| CityWarsaw | 490.092 | 26.931 | 0.000 |
| CFU_pres_abspos | 122.030 | 17.478 | 0.000 |
| CityCopenhagen:CFU_pres_abspos | 148.919 | 124.479 | 0.232 |
| CityRome:CFU_pres_abspos | -99.618 | 42.250 | 0.018 |
| CityWarsaw:CFU_pres_abspos | -364.220 | 30.186 | 0.000 |

Pairwise comparisons of each city pair was carried out by computing linear combinations of fixed effects with the betta_lincom function in the breakaway package. p-values were generated from a two-sided Wald test, against the null hypothesis that the linear combination of fixed effects is zero.

**Table S2.** Pairwise comparisons of city population richness estimates

| City pair | Estimates | Standard Errors | Lower CIs | Upper CIs | p-values | Adjusted p-value<sup>1</sup> |
| --- | --- | --- | --- | --- | --- | --- |
| Athens - Copenhagen | -352.295 | 77.950 | -505.074 | -199.516 | 3.1e-06 | 0.000 |
| Athens - Rome | 54.415 | 67.698 | -78.270 | 187.100 | 0.211 | 0.253 |
| Athens - Warsaw | -392.594 | 87.263 | -563.627 | -221.561 | 3.41e-06 | 0.000 |
| Copenhagen - Rome | 406.710 | 50.392 | 307.944 | 505.476 | 3.49e-16 | 0.000 |
| Copenhagen - Warsaw | -40.299 | 74.641 | -186.592 | 105.994 | 0.295 | 0.295 |
| Rome - Warsaw | -447.009 | 63.859 | -572.171 | -321.847 | 1.28e-12 | 0.000 |

<sup>1</sup>p-values corrected for multiple comparisons using the Benjamini & Hochberg method

#### Shannon index

The sample Shannon index was calculated from the normalised data set (n = 10 148 sequences per sample), using the phyloseq package, and a mixed effects model was built with Shannon index as the response variable. City and presence/absence of culturable *Legionella* were specified as fixed effects (including their interaction), and plumbing compartment, nested within building ID, were specified as random effects. Overall significance of the model fixed effects was tested using ANOVA.

**Table S3.** Sample Shannon index: ANOVA of mixed effect model. Model formula: Shannon ∼ City * CFU_pres_abs + (1 | Bldg_ID / Location_in_building)

|  | Sum Sq | Mean Sq | NumDF | DenDF | F value | Pr(>F) |
| --- | --- | --- | --- | --- | --- | --- |
| City | 6.166 | 2.055 | 3 | 9.991 | 2.732 | 0.100 |
| CFU_pres_abs | 2.765 | 2.765 | 1 | 135.410 | 3.674 | 0.057 |
| City:CFU_pres_abs | 2.347 | 0.782 | 3 | 127.919 | 1.040 | 0.377 |

Pairwise comparisons of city Shannon indices were carried out from the model output with the lsmeans package.

**Table S4.** Pairwise comparisons of city Shannon diversity indices

| City pair | estimate | SE | df | t.ratio | p.value<sup>1</sup> |
| --- | --- | --- | --- | --- | --- |
| Athens - Rome | 0.270 | 0.389 | 8.152 | 0.695 | 0.896 |
| Athens - Warsaw | -0.699 | 0.386 | 7.940 | -1.812 | 0.335 |
| Athens - Copenhagen | -0.559 | 0.426 | 10.832 | -1.312 | 0.575 |
| Rome - Warsaw | -0.969 | 0.383 | 8.307 | -2.529 | 0.127 |
| Rome - Copenhagen | -0.829 | 0.424 | 11.315 | -1.957 | 0.260 |
| Warsaw - Copenhagen | 0.140 | 0.421 | 11.081 | 0.332 | 0.987 |

<sup>1</sup>p-values corrected for multiple comparisons using the Tukey method

#### PERMANOVA on distance matrix (adonis)

For beta diversity analyses, we used the compositional Aitchison distance, equivalent to center-log-ratio transformation of the ASV table followed by Euclidean distance calculation. The influence of city and presence/absence of culturable Legionella (plus their interaction) was tested by permutational multivariate analysis of variance (PERMANOVA), using the adonis function in the vegan package.

**Table S5.** PERMANOVA of Aitchison distance matrix, computed using adonis function with strata = Building_loc_ID to account for repeated measures. Formula: adonis(clr_dist_matrix ∼ City * CFU_pres_abs, strata = Bldg_loc_ID, permutations = 10000)

|  | Df | SumsOfSqs | MeanSqs | F.Model | R2 | Pr(>F) |
| --- | --- | --- | --- | --- | --- | --- |
| City | 3 | 172 134 | 57 378 | 14.2983 | 0.2008 | 0.0014 |
| CFU_pres_abs | 1 | 22 919 | 22 919 | 5.7114 | 0.0267 | 0.0005 |
| City:CFU_pres_abs | 3 | 31 990 | 10 663 | 2.6573 | 0.0373 | 0.0545 |
| Residuals | 157 | 630 027 | 4 013 |  | 0.7351 |  |
| Total | 164 | 857 070 |  |  | 1.0000 |  |

#### Dispersion test (betadispr with permutest)

Multivariate homogeneity of group dispersions was tested for city and *Legionella* pres/abs with the betadisper function in the vegan package.

**Table S6.** Dispersion of groups around centroid, computed using betadispr function in vegan package

| Variable |  | Df | Sum Sq | Mean Sq | F | N.Perm | Pr(>F) |
| --- | --- | --- | --- | --- | --- | --- | --- |
| City | Groups | 3 | 12 405 | 4 135.01 | 15.3658 | 10 000 | 0.0001 |
|  | Residuals | 161 | 43 326 | 269.10 |  |  |  |
| CFU pres/abs | Groups | 1 | 40 | 39.89 | 0.0774 | 10 000 | 0.7764 |
|  | Residuals | 163 | 84 004 | 515.36 |  |  |  |

#### Culturable Legionella (CFU/L) by city

To test whether the Legionella culture counts (in CFU/L) differed among the cities, we build a generalised linear model to account for the Poisson distribution of counts. Model fit was improved by addition of random effects, and by separate modelling of zero inflation by city and random effects. Model construction steps are shown in Table S7, final model output in Table S8, and pairwise comparisons of cities in Table S9.

**Table S7.** Model selection for CFU/L data: GLM with Poisson distribution, with addition of zero inflation and mixed effects

| Model name | df | AIC | Counts model terms | Zeros counts model terms |
| --- | --- | --- | --- | --- |
| mod.poisson | 4 | 4 497 813 | ~ City | ~ 0 (no zero inflation modelling) |
| mod.poisson.zi.simple | 5 | 3 097 682 | ~ City | ~ 1 (constant zero modelling) |
| mod.poisson.zi.city | 8 | 3 097 647 | ~ City | ~ City |
| mod.poisson.me.zi.simple | 7 | 1 177 111 | ~ City +<br>(1\|Bldg_ID/Location_in_building) | ~ 1 (constant zero modelling) |
| mod.poisson.me.zi.city | 10 | 1 177 104 | ~ City +<br>(1\|Bldg_ID/Location_in_building) | ~ City |
| mod.poisson.me.zi.cityme | 12 | 1 177 090 | ~ City +<br>(1\|Bldg_ID/Location_in_building) | ~ City +<br>(1\|Bldg_ID/Location_in_building) |

**Table S8.** Culturable Legionella CFU/L by city: zero-inflated mixed effects generalised linear model with Poisson distribution (CFU/L counts and zero counts independently modelled by ∼ City + (1|Bldg_ID/Location_in_building)

| Model component |  | Estimate | Std. Error | z value | Pr(>\|z\|) |
| --- | --- | --- | --- | --- | --- |
| Counts | (Intercept) | -3.641 | 3.311 | -1.100 | 0.271 |
|  | CityAthens | 6.559 | 4.159 | 1.577 | 0.115 |
|  | CityRome | 7.610 | 4.075 | 1.868 | 0.062 |
|  | CityWarsaw | 12.690 | 4.198 | 3.023 | 0.003 |
| Zero counts | (Intercept) | 0.608 | 1.432 | 0.425 | 0.671 |
|  | CityAthens | -1.839 | 1.668 | -1.102 | 0.270 |
|  | CityRome | 0.624 | 1.697 | 0.368 | 0.713 |
|  | CityWarsaw | -2.146 | 1.725 | -1.244 | 0.214 |

**Table S9.** Pairwise comparisons of city culturable Legionella counts (CFU/L), computed from GLMM output using the lsmeans package.

| City pair | estimate | SE | df | t.ratio | p.value<sup>1</sup> |
| --- | --- | --- | --- | --- | --- |
| Copenhagen - Athens | -6.559 | 4.159 | 153.000 | -1.577 | 0.395 |
| Copenhagen - Rome | -7.610 | 4.075 | 153.000 | -1.868 | 0.246 |
| Copenhagen - Warsaw | -12.690 | 4.198 | 153.000 | -3.023 | 0.015 |
| Athens - Rome | -1.051 | 3.874 | 153.000 | -0.271 | 0.993 |
| Athens - Warsaw | -6.131 | 3.669 | 153.000 | -1.671 | 0.342 |
| Rome - Warsaw | -5.080 | 3.883 | 153.000 | -1.308 | 0.559 |

<sup>1</sup>p-values corrected for multiple comparisons using the Tukey method

#### Legionella genus in all 16S rRNA data

The proportions of all 16S rRNA sequences that classified as the genus Legionella, and as the species L. pneumophila, were calculated for each sample, as well as the proportion of Legionella genus sequences that were further classified as L. pneumophila. Mixed effects models were built with these proportions as response variables. City and presence/absence of culturable *Legionella* were specified as fixed effects (including their interaction), and plumbing compartment, nested within building ID, were specified as random effects. Overall significance of the model fixed effects was tested using ANOVA.

**Table S10.**
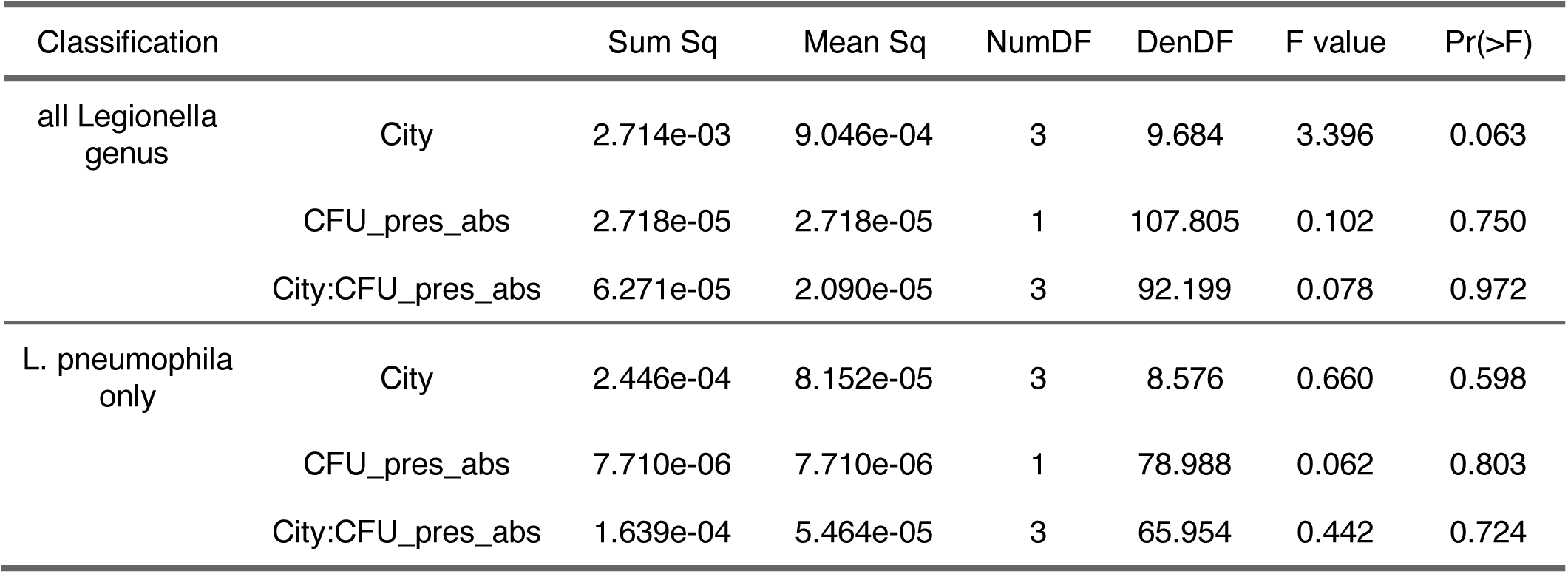
Proportion Legionella genus and L. pneumophila in 16S rRNA data: ANOVA of mixed effect models. Model formula: Proportion ∼ City * CFU_pres_abs + (1 | Bldg_ID / Location_in_building)

| Classification |  | Sum Sq | Mean Sq | NumDF | DenDF | F value | Pr(>F) |
| --- | --- | --- | --- | --- | --- | --- | --- |
| all *Legionella* genus | City | 2.714e-03 | 9.046e-04 | 3 | 9.684 | 3.396 | 0.063 |
|  | CFU_pres_abs | 2.718e-05 | 2.718e-05 | 1 | 107.805 | 0.102 | 0.750 |
|  | City:CFU_pres_abs | 6.271e-05 | 2.090e-05 | 3 | 92.199 | 0.078 | 0.972 |
| *L. pneumophila* only | City | 2.446e-04 | 8.152e-05 | 3 | 8.576 | 0.660 | 0.598 |
|  | CFU_pres_abs | 7.710e-06 | 7.710e-06 | 1 | 78.988 | 0.062 | 0.803 |
|  | City:CFU_pres_abs | 1.639e-04 | 5.464e-05 | 3 | 65.954 | 0.442 | 0.724 |

**Table S11.** Proportion of all Legionella genus 16S sequences that are further classified as L. pneumophila: ANOVA of mixed effect model. Model formula: L.pneumo proportion ∼ City * CFU_pres_abs + (1 | Bldg_ID / Location_in_building)

|  | Sum Sq | Mean Sq | NumDF | DenDF | F value | Pr(>F) |
| --- | --- | --- | --- | --- | --- | --- |
| City | 4 182.162 | 1 394.054 | 3 | 8.032 | 1.858 | 0.215 |
| CFU_pres_abs | 5 271.805 | 5 271.805 | 1 | 135.533 | 7.027 | 0.009 |
| City:CFU_pres_abs | 4 761.232 | 1 587.077 | 3 | 115.020 | 2.116 | 0.102 |

